# Sexual Dimorphic Gene Expression Profile of Perirenal Adipose Tissue in Ovine Fetuses with Growth Restriction

**DOI:** 10.1101/2023.03.04.531079

**Authors:** Jack Blomberg, Rosa I. Luna Ramirez, Dipali Goyal, Sean W. Limesand, Ravi Goyal

**Author notes:** **Correspondence:** Ravi Goyal, MD, PhD, Associate Professor, School of Animal and Comparative Biomedical Sciences, College of Agriculture and Life Sciences, The University of Arizona, Tucson, AZ 85719.

## Abstract

Worldwide, fetal growth restriction (FGR) affects 7 to 10% of pregnancies, or roughly 20.5 million infants, each year. FGR not only increases neonatal mortality and morbidity but also the risk of obesity in later life. Currently, the molecular mechanisms by which FGR “programs” an obese phenotype are not well understood. Studies demonstrate that FGR females are more prone to obesity compared to males; however, the molecular mechanisms that lead to the sexually dimorphic programming of FGR are not known. Thus, *we hypothesized that FGR leads to the sexually dimorphic programming of preadipocytes and reduces their ability to differentiate into mature adipocytes*. To test the hypothesis, we utilized a maternal hyperthermia-induced placental insufficiency to restrict fetal growth in sheep. We collected perirenal adipose tissue from male and female near-term FGR and normal-weight fetal lambs (N=4 in each group, 16 total), examined the preadipocytes’ differentiation potential, and identified differential mRNA transcript expression in perirenal adipose tissue. Male FGR fetuses have lower cellular density compared to control male fetuses. However, no difference was observed in female FGR fetuses compared to control female fetuses. In addition, the ability of preadipocytes to differentiate into mature adipocytes with fat accumulation was impaired in male FGR fetuses, but this was not observed in female FGR fetuses. Finally, we examined the genes and pathways involved in the sexually dimorphic programming of obesity by FGR. On enrichment of differentially expressed genes in males compared to females, the Thermogenesis KEGG Pathway was downregulated, and the Metabolic and Steroid Biosynthesis KEGG pathways were upregulated. On enrichment of differentially expressed genes in male FGR compared to male control, the Steroid Biosynthesis KEGG Pathway was downregulated, and the PPAR Signaling KEGG pathway was upregulated. No pathways were altered in females in response to growth restriction in perirenal adipose tissue. Thus, the present study demonstrates a sexually dimorphic program in response to growth restriction in sheep fetal perirenal adipose tissue.

## Introduction

Obesity has emerged as a major health crisis in the USA and worldwide. According to the WHO, the prevalence of obesity has tripled since 1975 (1). Based on the CDC data, the prevalence of obesity was 42.4% in 2017 and 2018. Financially, obesity plays a large part in higher medical costs, decreased productivity, and increased absence from work (2). With healthcare costs rising every day, the cost of excess healthcare expenditures takes an enormous amount of toll on individuals. In 2014, globally, the economic impact of obesity was $2.0 trillion US dollars (2). Additionally, obesity significantly reduces life span and quality of life. It is shown that obesity can decrease life expectancy by as much as 5-10 years (3). Obesity is related to disorders such as coronary artery disease, arthritis, and stroke. Obesity also has a considerable impact on mental health conditions like low self-esteem, mood disorder, and eating disorders (4). Overall, obesity is immensely impactful on society and requires urgent attention and investigation.

Human epidemiological studies demonstrate that FGR is a major risk factor for obesity later in life (5). This has led to the concept of <u>developmental origins of health and disease</u> (DOHaD) and “fetal programming” of obesity with associated metabolic disorders (6). Several studies support that FGR predisposes individuals to a significantly higher risk of developing obesity (7, 8). The thrifty phenotype hypothesis explains the mechanism of fetal programming that the in-utero environment of reduced nutrient availability prepares the fetus for the predicted outside environment of nutrient scarcity after birth (6). Several studies support that a fetus is “programmed” to live under thrifty conditions (7, 9–13). If, postnatally, there is adequate nutrition available to the newborn, it starts accumulating excess nutrition as fat for future nutritional starvation. This leads to rapid catch-up growth, which is a risk factor for metabolic syndrome (7, 14). Mechanisms of such programming are not entirely understood (9, 10). Nonetheless, it is well established that FGR predisposes to obese phenotype (9, 15–17). Obesity is a very complex process and depends on the precise regulation of the gene expression (18). This complexity is in part because of the complex composition of adipose tissue, which contains several different cell types such as white adipocytes, which can accumulate fat, and brown adipocytes, which efficiently convert energy into heat, an intermediate phenotype known as beige adipocytes, and cells with stem cells like property (known as preadipocytes, vascular stromal fraction, or adipose tissue-derived stem cells). These preadipocytes are very plastic cell types (19). For instance, it can be differentiated into all three lineages - endodermal (20), mesodermal (21), and ectodermal origins (22). However, FGR-mediated programming of adipocytes is not well investigated and creates a significant knowledge gap in our current understanding. In the mouse, we have demonstrated that FGR pups are more prone to obesity and hypertension in a sexually dimorphic manner (7). Others have also demonstrated that FGR babies are at significant risk for excessive fat deposition in adipose tissues and rapid catch-up growth in a sexually dimorphic manner (6, 23, 24). Specifically, FGR females are prone to develop greater obesity compared to males in later life. Thus, *we hypothesized that FGR leads to the sexually dimorphic programming of preadipocytes and reduces their ability to differentiate into mature adipocytes*.

## Methods

All animal studies were reviewed and approved by the Institutional Animal Care and Use Committee (IACUC) of the University of Arizona. Crossbred Columbia-Rambouillet ewes carrying singleton pregnancies were used for the present study. All studies were conducted on the four study groups – near-term control male fetuses (MC), near-term control female fetuses (FC), near-term FGR male fetuses (MFGR), and near-term FGR female fetuses (FFGR).

### FGR Animal Model

Pregnant ewes were exposed to high ambient temperatures to produce maternal hyperthermia that causes progressive placental insufficiency and fetal growth restriction (25, 26). In this model of FGR, pregnant ewes are exposed to elevated ambient temperatures (40°C for 12 hours; 35°C for 12 hours; dew point 22°C) from 38 ± 1 to 87 ± 1 days of gestation (total gestation in sheep is ~149 days). Control fetuses are from ewes maintained at 22 ± 1°C and pair-fed to the average *ad libitum* feed intake of the hyperthermic group (25, 27–31). All ewes are given *ad libitum* access to water and salt. Following physiological measurements previously reported (32), the ewe and fetus were killed humanely (Euthasol; Virbac Animal Health).

### Adipose Tissue Collection

Perirenal adipose tissue from four experimental groups was collected to examine morphological differences and changes in gene expression. Perirenal adipose tissue is a visceral subtype with well-defined boundaries and is covered with a perirenal fascia (Gerota’s fascia), which allows for complete dissection and weight measurements. After weighing the adipose tissue, a portion of perirenal adipose tissue was used to isolate preadipocytes, a slice was fixed in 4% paraformaldehyde, and the rest was stored in liquid nitrogen for downstream RNAseq analysis (29, 33).

### Cell Density in Adipose Tissue

Cell Density was examined by staining the section with Hoechst live-cell stains. The blue nuclei stained were counted using ImageJ software.

### Preadipocyte Isolation

Preadipocytes were isolated using a published protocol (34). Briefly, 500 mg of adipose tissue was minced in a sterile Petri dish, and 2 ml of 5 mg/ml collagenase in sterile PBS was added to the minced tube and incubated in an orbital shaker for 1 hour at 37°C at 400 RPM. The tissue was triturated after every 15 minutes. Following the incubation, the media at the bottom of the tube was removed and filtered with a 40-micron filter. The filtrate was centrifuged at 1000 g for 5 minutes. The pellet was mixed in RBC lysis buffer (Fisher Scientific, Cat # 501129743) and incubated at room temperature for 10 minutes. The cells were re-pelleted by centrifugation at 500 g for 5 minutes. The pelleted cells were mixed in DMEM with 10% FBS and 1% Pen-Strep to the culture at 37 C in 5% CO2 and air. The cells were washed the next day with sterile PBS, and the cells which adhered to the plastic surface of the cell culture flask were passaged further.

### Preadipocyte Differentiation

Adipogenic differentiation was initiated by culturing the preadipocytes in adipogenesis initiation media (DMEM with 10% FBS, 1 μM insulin, 1 μM Dexamethasone, and 500 μM 3-isobutyl-1-methylxanthine (IBMX) and 1% penicillinstreptomycin) for 7 days. Following this, adipogenesis differentiation media (DMEM with 10% FBS, 1 μM insulin, 1 μM Dexamethasone, indomethacin 50 μM, rosiglitazone 20 μM, and 500 μM IBMX was added for the next 9 days. After 16 days of differentiation, Oil Red Staining was conducted by fixing the cells in calcium fixative and oil red staining solution as published (35).

### Whole Transcriptomic Analysis

To determine the mechanistic pathways involved in this sexually dimorphic programming of adipocytes, we conducted RNAseq on the perirenal adipose tissue from the four groups following standard protocol. Briefly, the stored adipose tissue was thawed in Trizol and homogenized using 3 pulses, 4-6 seconds each. Then, 0.2 mL of chloroform was added per 1 mL of Trizol (0.1 mL per sample). Samples were centrifuged for 15 minutes at 14,000 x g (4 °C) and the mixture separated into a lower red phenol-chloroform, interphase, and a colorless upper aqueous solution. The aqueous phase containing RNA was transferred to a new tube very carefully to ensure proteins and other cell components did not contaminate the RNA. Then, 250 μl of isopropanol was added to the aqueous phase (0.5 ml/ ml of Trizol) and incubated for 10 minutes. The sample was centrifuged for 10 minutes at 14,000 x g (4°C) and the RNA precipitate formed a white gel-like pellet at the bottom of the tube. The supernatant was discarded and the sample was vortexed briefly before centrifugation for 5 minutes at 8000 x g (4°C). The supernatant was discarded and the RNA pellet was air-dried for 10 minutes. Pellets were resuspended in 50 μl of RNase-free water. The RNA was further purified by Zymo Purelink RNA columns. The obtained RNA samples were measured for quantity (ng/ml) and purity on Nanodrop and Qubit (broad-range) according to 260/280 nucleic acid absorbance ratio before sending the University of Arizona Genetics Core Facility for sequencing, where RNA were further checked for quality and quantity with an Advanced Analytics Fragment Analyzer (*High Sensitivity RNA Analysis Kit – Catalog # DNF-491*) and quantity with determined with a Qubit RNA quantification kit (*Qubit® RNA HS Assay Kit – Catalog # Q32852*).

Once quality and quantity were validated, a library was constructed from samples using a Swift RNA Library Kit – (*Catalog # R1024*) and Swift Dual Combinatorial Indexing Kit – (*Catalog # X8096*). Upon constructing the library, the average fragment size in the library was determined with the Advanced Analytics Fragment Analyzer with the High Sensitivity NGS Analysis Kit *– (Catalog # DNF-486*). Quantity was evaluated with an Illumina Universal Adaptor-specific qPCR kit, the Kapa Library Quantification Kit for Illumina NGS *– (Catalog # KK4824*).

After completing the final library QC, samples were equimolar-pooled and clustered for sequencing on the NextSeq500 machine. The sequencing run was performed using Illumina NextSeq500 run chemistry (*NextSeq 500/550 High Output v2 kit 150 cycles – Catalog FC-404-2002*).

#### RNAseq Data Analysis

Sequence data quality was validated for RNAseq analysis using FastQC Version 0.11.9. Sequences with average Phred scores below 34 were discarded. Fully annotated genome indices were generated for sheep (Oar_rambouillet_v1.0), cow (ARS-UCD1.2), mouse (GRCm39), and human (GRCh38.p13) using ENSEMBL (v1.0.103) and aligned with the sequencing data using Salmon (36). Integrated differential expression and pathway analysis were conducted with iDep.96 web-based applications (37). During preprocessing, the genes with less than 20 counts per million in 3 or more libraries were discarded from further processing, and counts were normalized using EDGER: log2(CPM+c) transformation. The distribution of the transformed data box and density plots are provided in Supplemental Figure 1. Enrichment analysis for differentially expressed genes was conducted using a False Discovery Rate (FDR) cutoff of 0.05 and a minimum fold change of 1.5. Genes altered were matched with the known Kyoto encyclopedia of genes and genomes (KEGG) pathways (38), and gene set enrichment analysis (GSEA) was conducted as published (39). Generally Applicable Gene-set Enrichment (GAGE) was also conducted to determine the biological relevance of the regulatory mechanism (40).

**Figure 1:**
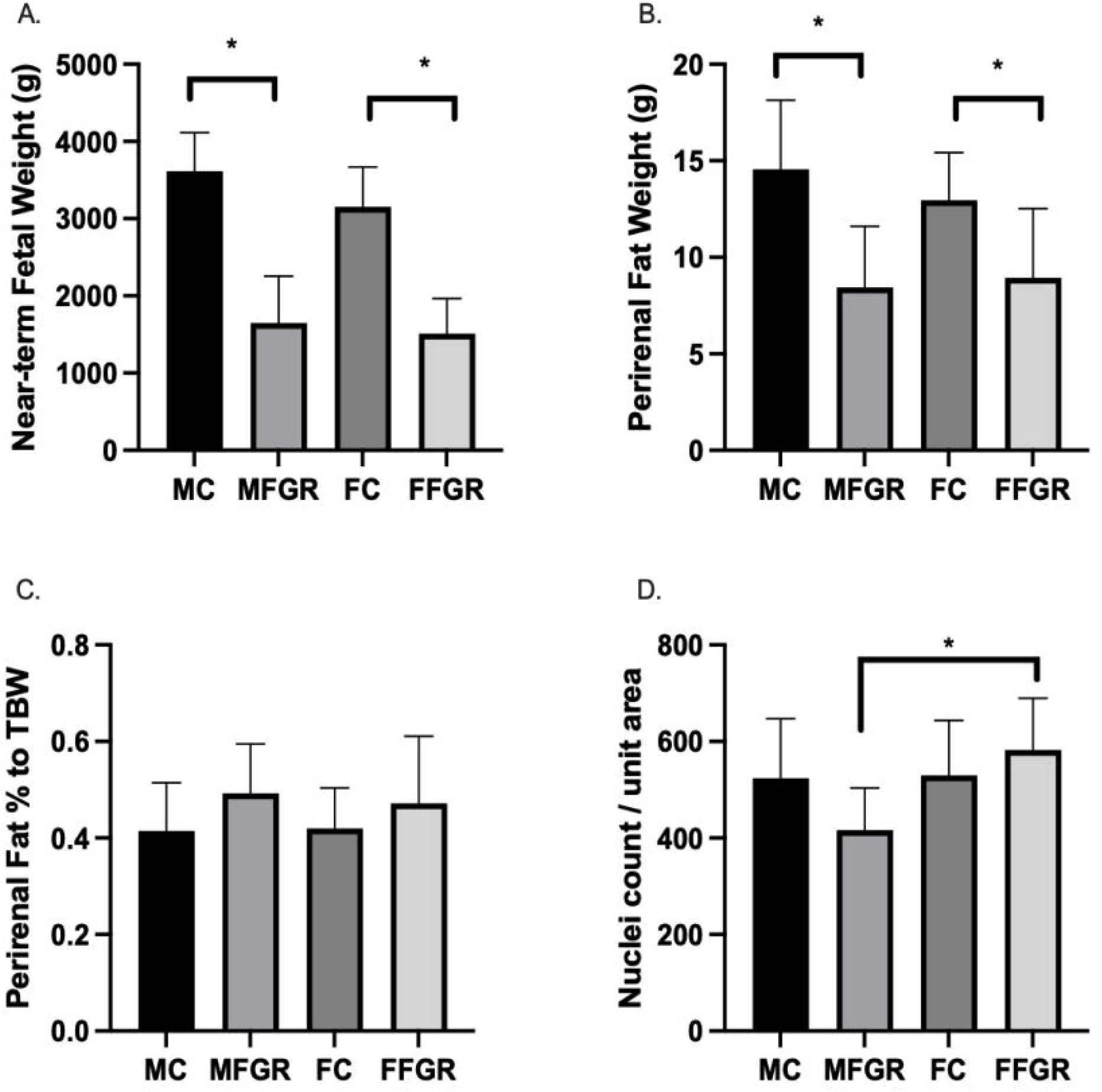
Bar graphs demonstrate: (A) Body weights (B) Perirenal fat weights (C) Perirenal fat weight as a % of total body weight, and (D) Nuclei count per unit area of perirenal fat 10-micron slices from control and growth-restricted ovine near-term fetuses. N = 5 in each group and * denotes P < 0.05. MC = Male control, FC = Female Control, MFGR = Growth Restricted Male Fetus, and FFGR = Growth Restricted Female Fetus.

## Results

### Effect of Ambient Hyperthermia on Fetal and Perirenal Adipose Weight

Maternal hyperthermia reduced the body weight of both male and female fetuses (Figure 1A). The average perirenal adipose tissue weight was less in both FGR males and females compared to the control counterparts (Figure 1B). However, when perirenal adipose tissue was normalized to total fetal body weight (TBW), there was no difference between FGR and control groups in both sexes (Figure 1C).

### Cellular Density of Perirenal Adipose Tissue

The results demonstrate a significantly lower nuclei number/unit area in FGR males compared to all other groups (Figure 1D).

#### Sexually Dimorphic Programming of Preadipocyte Differentiation

The results (Figure 2) demonstrate that the preadipocyte differentiation into mature adipocytes containing lipid droplets as stained by oil red stain was significantly reduced in MFGR compared to all other groups. There was no significant effect of sex on the differentiation ability of preadipocytes into mature adipocytes in control males and females.

**Figure 2:**
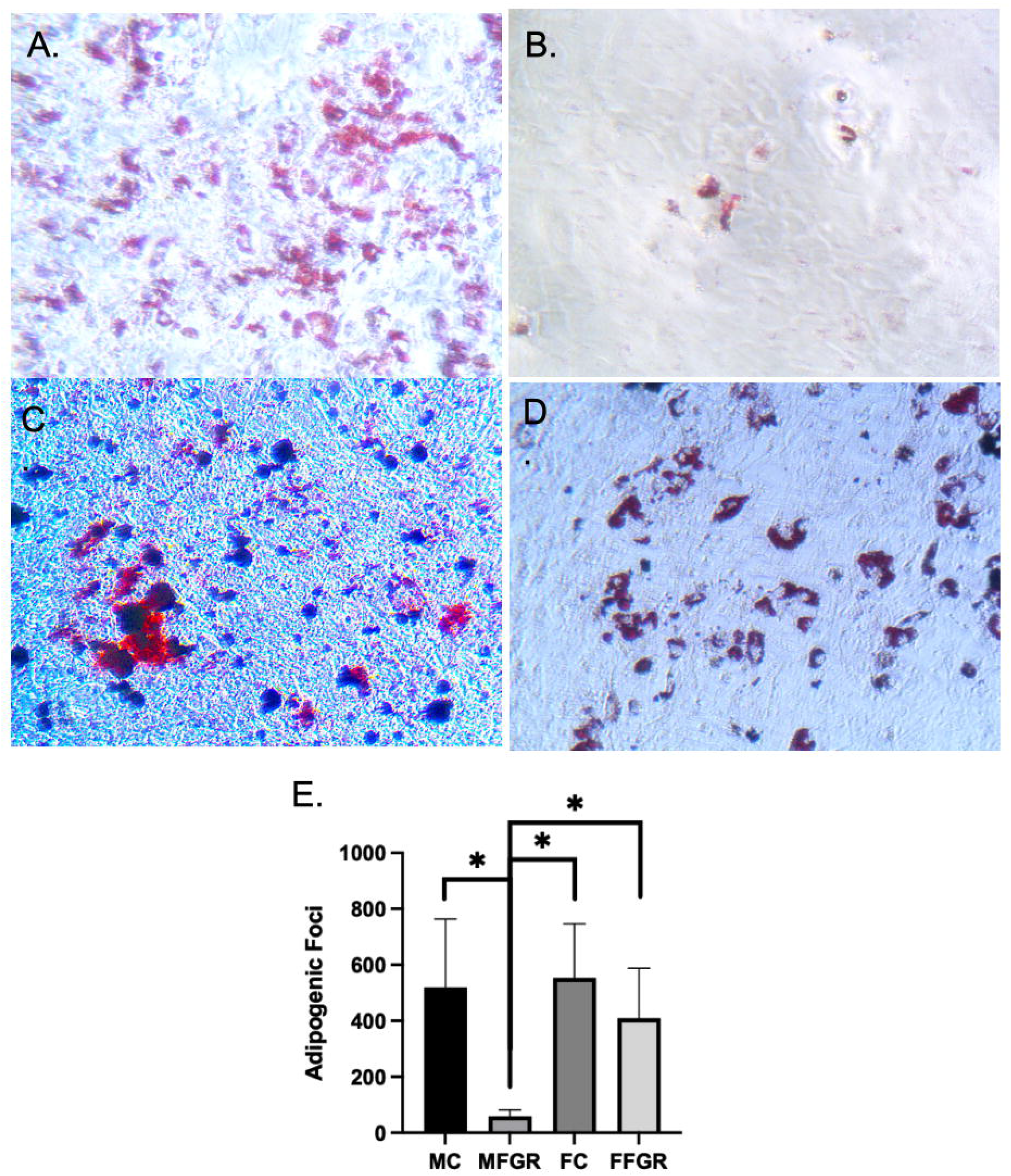
Demonstrates oil red staining of fat droplets following adipogenic differentiation of preadipocytes. (A) MC group. (B) MFGR group. (C) FC group. (D) FFGR group. (E) Bar graph of droplet counts. N = 5. * Denotes P < 0.05.

### RNAseq analysis of the perirenal adipose tissue

Results demonstrate a strong effect of sex on gene clustering (Figure 3A). The clustering analysis demonstrates that growth restriction did not have much effect on gene expression in female fetuses (Figure 3B). In contrast, in male fetuses, the gene groups clustered separately following growth restriction (Figure 3B). There were few differentially expressed genes in females compared to males with growth restriction (Figures 3C and D). A total of 1330 gene transcripts were differentially regulated (FDR < 0.05 and FC > 1.5) in male and female perirenal adipose tissue (Figure 3). Of these, 643 were upregulated, and 687 were downregulated in female fetus perirenal adipose tissue as compared to adipose tissue from male fetuses (Figures 4A-D). Similarly, 148 genes were upregulated, and 124 were downregulated in MFGR as compared to MC (Figures 4E-H). Surprisingly, in females, only 6 genes were upregulated and 2 downregulated in FFGR perirenal adipose tissue as compared to FC (Figures 4I-L). On comparative analysis, there was only 1 gene commonly altered in response to growth restriction in visceral adipose from both males and females (Novel gene ENSOARG00020003728). The complete list of differentially regulated genes in males as compared to females is provided in supplemental Tables S1, S2, and S3. The heat maps and principal component analysis demonstrate a separate clustering of males and females.

**Figure 3:**
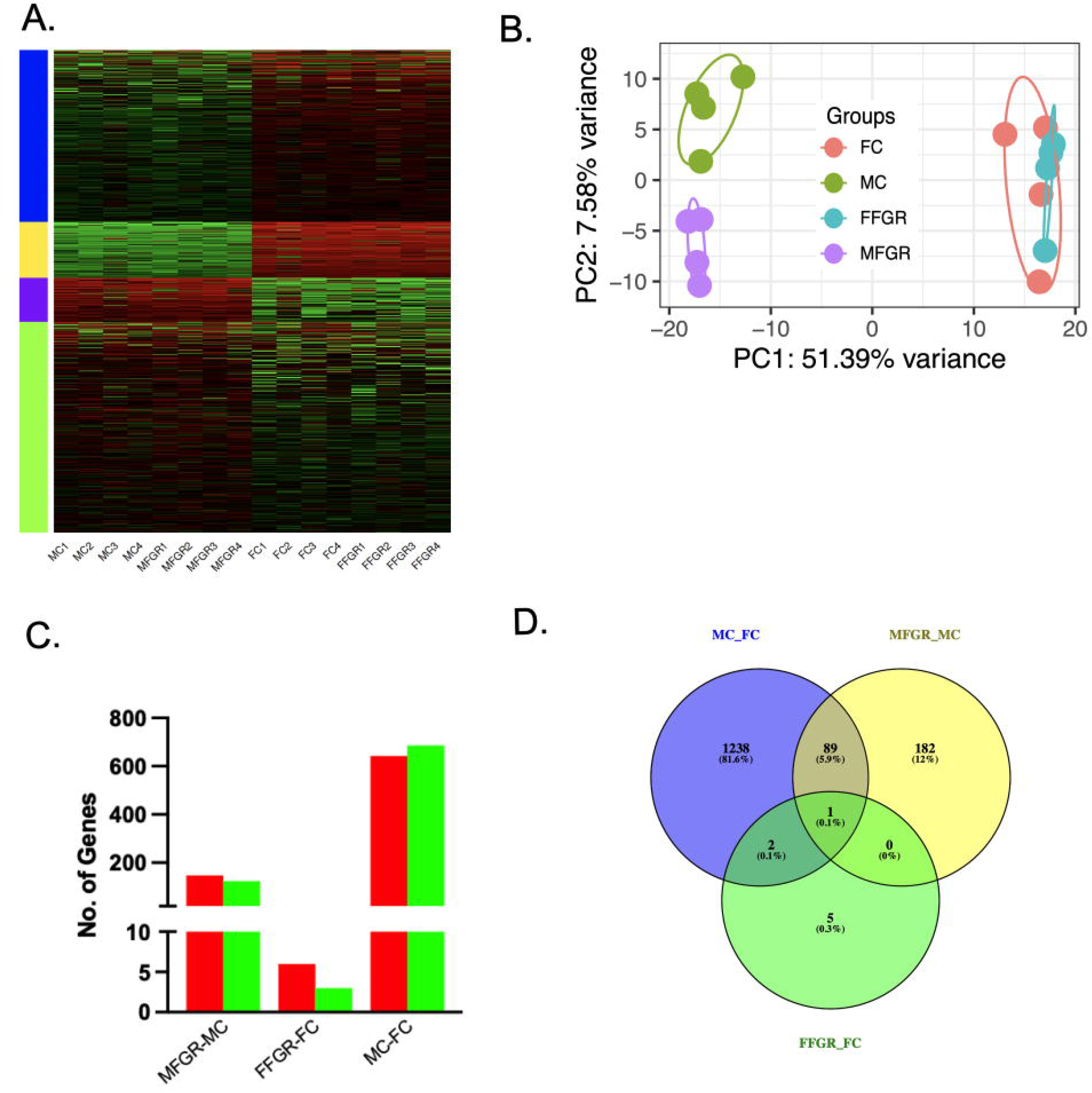
Sexually dimorphic clustering of gene expression in the four groups. (A) Hierarchical clustering of the differentially expressed genes in perirenal adipose tissue from male and female sheep fetuses. (B) PCA plot of the differential gene expression in the four study groups. (C) The bar graph demonstrates upregulated (red) and downregulated (green) genes by comparing the MFGR with MC, FFGR with FC, and MC with FC. (D) Venn diagram demonstrates the number of genes that overlapped with sex and growth restriction variables.

**Figure 4.**
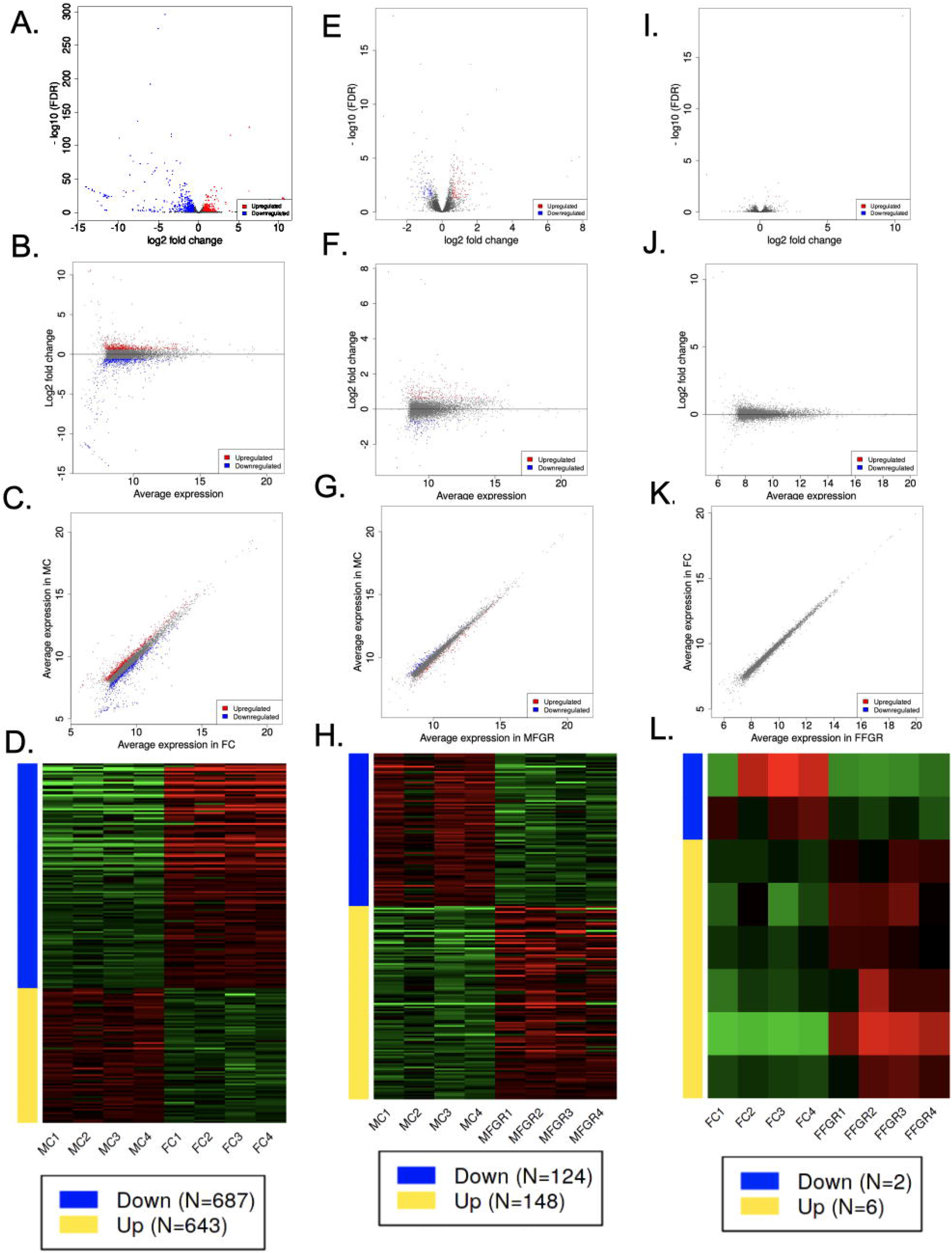
Demonstrates differential gene expression in the three comparisons (MC vs. FC; MFGR vs. MC; and FFGR vs FC). Panels A, E, and I are the volcano plots demonstrating -log10 FDR (Y-axis) and log2 fold change (X-axis) of the differentially expressed genes in the three comparative groups. Panels B, F, and J demonstrate the scatter plot of log2 fold change (Y-axis) with average expression (X-axis) of the differentially expressed genes in the three comparative groups. Panels C, G, and K demonstrate the scatter plots comparing the average expression of differentially expressed genes in the members of each group. Panels D, H, and L demonstrate the hierarchical clustering of the differentially regulated genes in each comparative group.

### Gene Enrichment and Pathway Analysis

#### Effect of sex on visceral adipose tissue gene expression pathways

We observed a significant effect of sex on gene expression in visceral adipose tissue (Figures 3 & 4). On enrichment of the genes altered in males compared to females, the major network upregulated in males were those involved in neovascularization and downregulated network involved mitochondrial function (Figure 5). On enrichment of differentially expressed genes in males compared to females, the Thermogenesis KEGG Pathway was downregulated, and the Metabolic and Steroid Biosynthesis KEGG pathways were upregulated (Supplemental Table S4).

**Figure 5.**
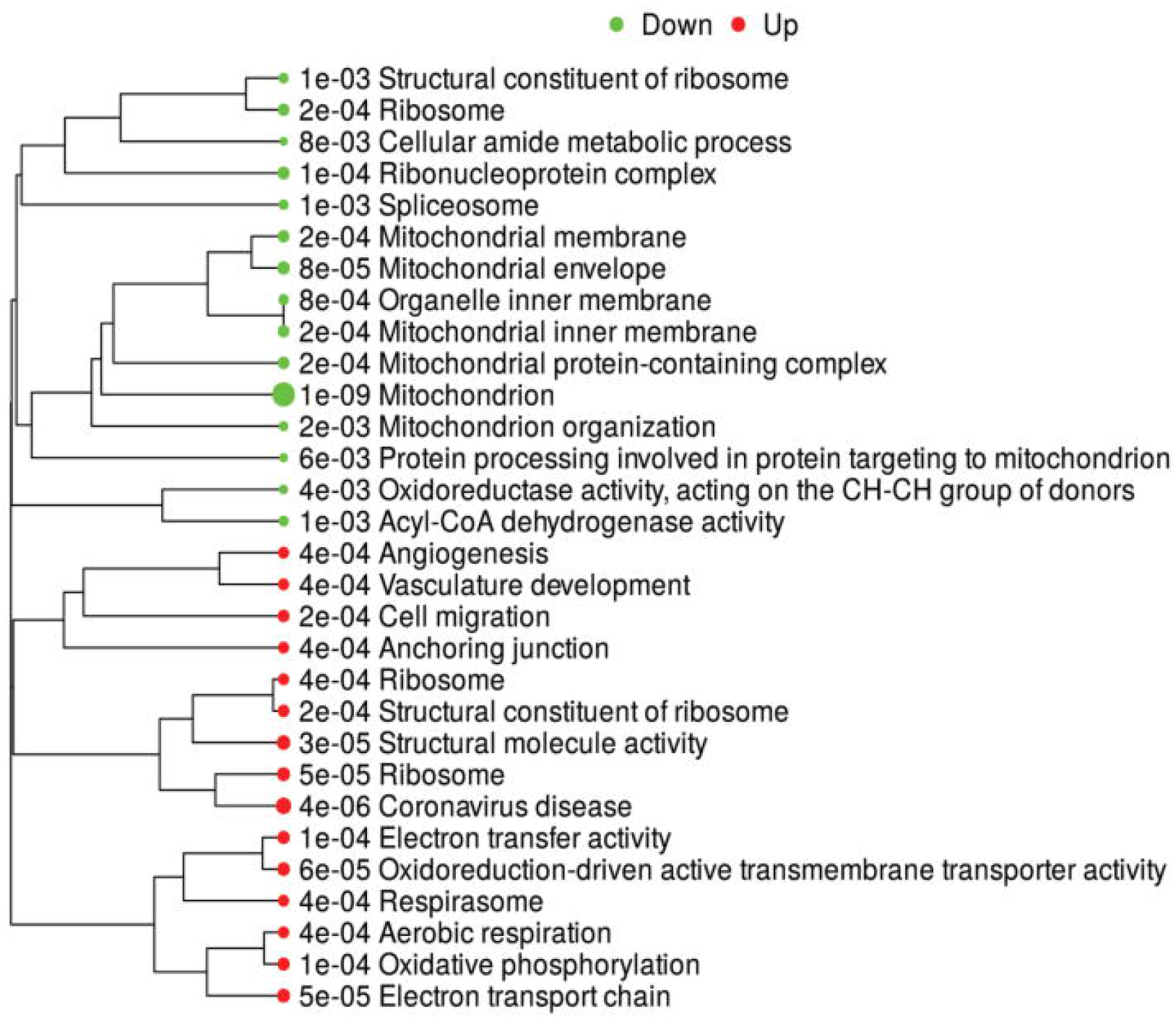
Demonstrates enriched pathways in DEGs for genes altered on comparing male vs. female control fetuses. Green dots indicate downregulated pathways, and red dots indicate upregulated pathways. The sizes of the dots correspond to the adjusted P-value.

**Figure 6.**
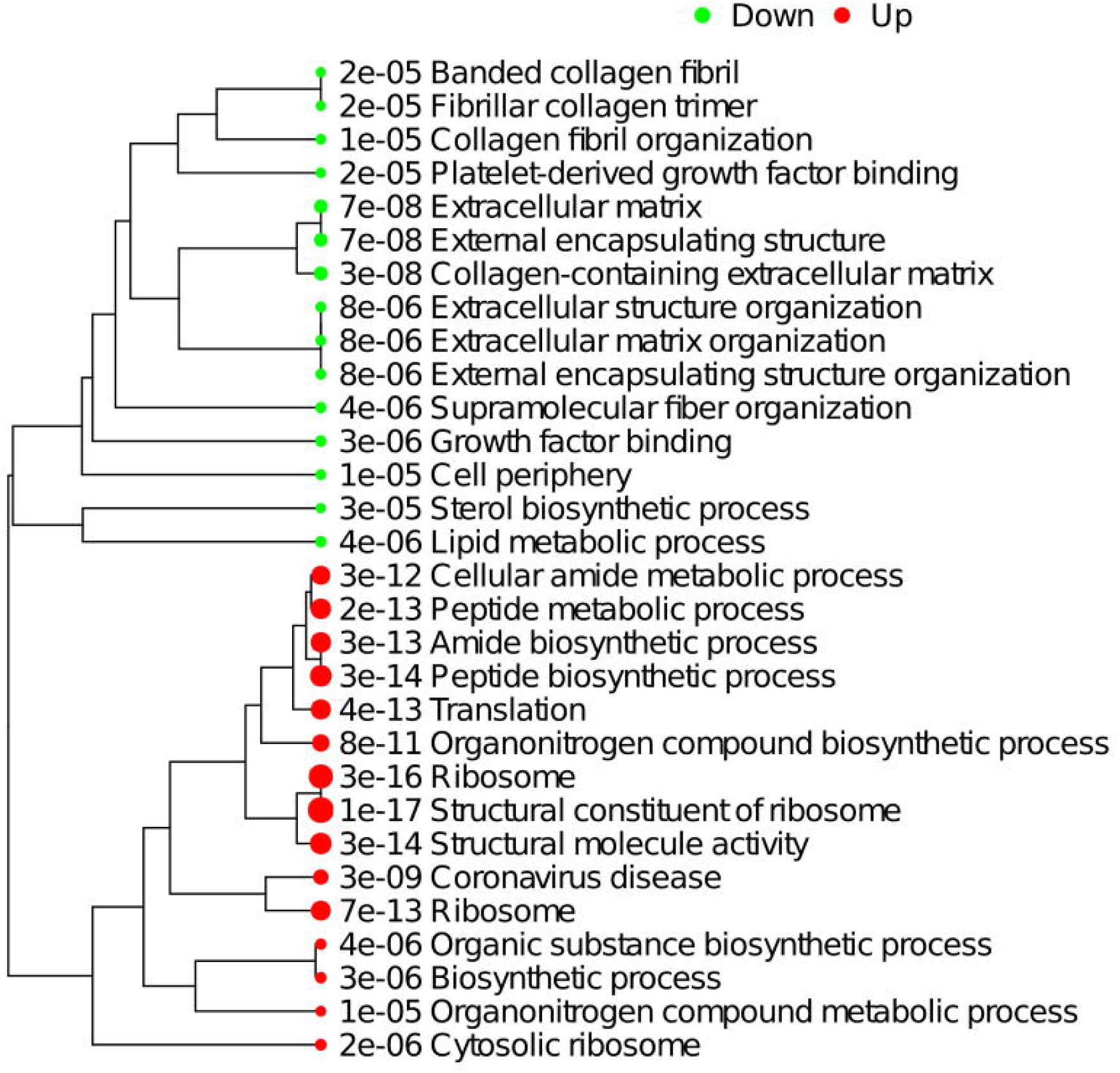
Demonstrates enriched pathways in DEGs for genes altered on comparing MFGR vs. MC. Green dots indicate downregulated pathways, and red dots indicate upregulated pathways. The sizes of the dots correspond to the adjusted P-value.

#### Effect of growth restriction on gene expression pathways

The result demonstrates that female adipose tissue did not undergo major changes with growth restriction. Only 8 genes were altered in FGR females compared to control females, and they did not map to a common pathway. However, following fetal growth restriction, the translation and biosynthetic pathways were upregulated in males, and lipid/sterol pathways were downregulated. On enrichment of differentially expressed genes in MFGR compared to MC, the Steroid Biosynthesis KEGG Pathway was downregulated, and the PPAR Signaling KEGG pathway was upregulated.

### Relative expression of preadipocytes, mature adipocytes, white adipocytes, and brown adipocytes markers in the four study groups

THY1 was used as preadipocyte markers; ADIPOQ and FABP4 were used as mature adipocyte markers, UCP1 and CIDEA were used as brown adipose markers; and TCF21 and leptin were used as white adipose tissue markers. Of note, leptin is also considered a mature adipose tissue marker. There was no change in any markers in females with growth restriction. In males, FABP4, a mature adipocyte marker, was significantly upregulated (> 2.5 folds), and THY1, a preadipocyte marker, was downregulated (<0.5 fold). On examining the effect of sex on the markers, THY1 (preadipocyte marker) was upregulated in control males as compared to control females (>2 fold).

## Discussion

Fetal growth restriction is a known risk factor for obesity in adult life. The mechanisms of fetal programming of adult disease are still not well understood. In the present study, we demonstrate that ambient hyperthermia induces a significant growth restriction in fetuses. Although the overall perirenal weight was lower in growth-restricted fetuses, there was no significant difference with control fetuses following normalization to body weight. Several studies have shown that adipose tissue relative percentages are maintained or may be higher in an organism despite growth restriction (42–44). In contrast, several studies have demonstrated that the percentage weight of several organs relative to the total body weight is significantly lower in growth-restricted fetuses (45, 46). Notably, similar to adipose tissue, the brain is the other organ that shows relative sparing in weight reduction in response to growth restriction. Thus, active programming of the adipose tissue and protection indicates the importance of adipose tissue in organismal biology.

In recent years, adipose tissue has emerged as an important endocrine organ and is responsible for the secretion of several hormones (47). In the present report, irrespective of growth restriction, we observed significant differences in gene expression in male versus female perirenal adipose tissue. In males, the neovascularization network was upregulated as compared to that in females. Similarly, the mitochondria-related genes were downregulated in males as compared to those in adipose tissue from female fetuses. In a recent report, it was demonstrated that with a high-fat diet (HFD), there was an increase in mitochondrial complexes in female mice as compared to those in male mice, and the male mice were more prone to obesity (48). Similarly, sexual dimorphism in adipose tissue has been demonstrated by others in response to a high-fat diet in mice (49) and baboons (50). Thus, the present study further demonstrates sexually dimorphic programming in response to growth restriction in sheep adipose tissue. Also, in control males, we observed that the genes involved in thermogenesis pathways are downregulated, and those involved in metabolic pathways, including steroid biosynthesis, are upregulated. Reduction in thermogenesis may be responsible for making FGR males more prone to obesity than control.

Also, fetal growth restriction has been implicated in the “programming” of obesity in later life (9, 10). In the present report, we observed sexually dimorphic alterations in the differentiation ability of preadipocytes with growth restriction. There was a significant reduction in cellular density only in male growth-restricted fetuses. In growth-restricted male fetuses, there was also a significant reduction in the preadipocyte differentiation potential to mature adipocytes compared to male controls. These changes were not observed in female fetuses.

Similarly, in response to growth restriction, there was a significant change in the male transcriptome of perirenal adipose tissue, which was not observed in growth-restricted female fetuses. In growth-restricted male fetuses, the PPAR-gamma pathway was upregulated, and the PI3K-AKT pathway was downregulated with an associated increase in mature adipocyte marker and reduction in preadipocyte marker. Evidence supports that the PPAR-gamma pathway is involved in preadipocyte differentiation and maturation (51). Thus, it is possible that with growth restriction, male fetuses have a reduction in the preadipocyte population which may be the reason for reduced cell density in growth-restricted males.

A surprising finding of the present study was that transcriptomic profiles were unaffected by growth restriction in adipose tissue from female fetuses. In contrast, differences in male fetuses were observed at both transcriptomic and phenotypic levels. The significance of this sexually dimorphic programming and causal mechanisms warrants further investigation.

### Conclusion and Perspective

The present study demonstrates sexual dimorphic programming of perirenal adipose tissue differentiation and gene expression. Our findings show that female fetuses are more resistant than male fetuses. Following growth restriction, the adipose tissue from male fetuses had lower cell density, upregulated mature adipocyte markers, and lower in-vitro differentiation potential. Further studies are needed to examine if lower preadipocyte differentiation potential in males lead to more fat deposition from early life. The study also demonstrates that the PI3K-AKT and PPAR-gamma pathways require further investigation and may be involved in growth restriction-induced fetal programming of adipose tissue.

