## Supplementary figures and images for "Sexual Dimorphic Gene Expression Profile of Perirenal Adipose Tissue in Ovine Fetuses with Growth Restriction"

### Supplemental Figure 1

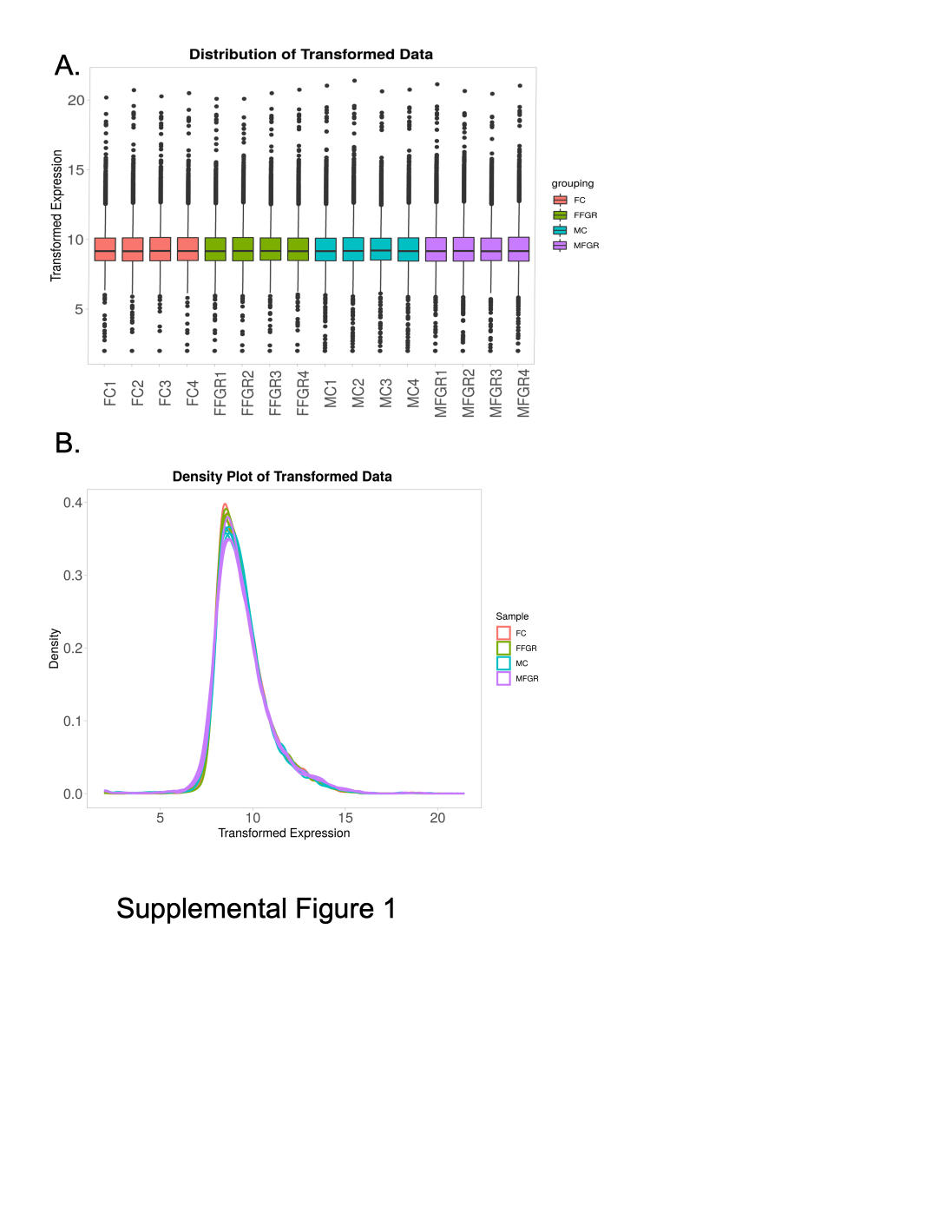
